# RIM-BP2 primes synaptic vesicles *via* recruitment of Munc13-1 at hippocampal mossy fiber synapses

**DOI:** 10.1101/469494

**Authors:** Marisa M Brockmann, Marta Maglione, Claudia G Willmes, Alexander Stumpf, A Boris Bouazza, Laura M Velasquez, Marie Katharina Grauel, Prateep Beed, J Stephan Sigrist, Christian Rosenmund, Dietmar Schmitz

## Abstract

All synapses require fusion-competent vesicles and coordinated Ca^2+^-secretion coupling for neurotransmission, yet functional and anatomical properties show a high diversity across different synapse types. We show here that the presynaptic protein RIM-BP2 has diversified functions in neurotransmitter release at different central mammalian synapses and thus contributes to synaptic diversity. At hippocampal pyramidal CA3-CA1 synapses, RIM-BP2 loss has a mild effect on neurotransmitter release, by only regulating Ca^2+^-secretion coupling. However, at hippocampal mossy fiber synapses RIM-BP2 has a strong impact on neurotransmitter release by promoting vesicle docking/priming *via* recruitment of Munc13-1. In wild type mossy fiber synapses, the distance between RIM-BP2 clusters and Munc13-1 clusters is larger than in hippocampal pyramidal CA3-CA1 synapses, suggesting that spatial organization may dictate the role a protein plays in synaptic transmission and that differences in active zone architecture is a major determinant factor in the functional diversity of synapses.

## Introduction

Across all types of synapses, vesicle fusion is coordinated by an evolutionarily conserved set of vesicular and active zone proteins^1^. One hallmark of synapses is their functional heterogeneity: indeed, synapses can exhibit high or low transmission fidelity and this diversity results in synapse-specific differences in response fluctuation and short-term plasticity^2,3^. In recent years, functional synaptic diversity has been found to be critical for routing and encoding sensory information within networks of neurons in the brain^4^. Functional synaptic diversity has been observed both within and across brain regions, and it has been shown to play a major role in temporal coding of multisensory integration and extraction of specific sensory features^1–4^. Still, the molecular origin of this heterogeneity is largely unknown, and analysis of genotype-phenotype differences across species and brain tissue have just started to uncover key molecular principles responsible for synaptic diversity, emphasizing the importance of abundance and isoforms differences across species for synaptic diversity^5–8^.

It is possible, but largely untested whether the active zone architecture, which is specialized throughout synapse types, is associated with distinct protein functions and thus contributes to synaptic diversity. Here, RIM-binding proteins (RIM-BPs) are particularly interesting, as their loss manifests in severe phenotypes in the drosophila neuromuscular junction (NMJ)^9^, but rather subtle phenotypes in small central murine synapses, the Calyx of Held or the ribbon synapse with only mild impairments in Ca^2+^-channel-release site coupling^10–12^. Drosophila NMJ and small central synapses are considerable distinct in their anatomical, ultrastructural and, physiological properties^13^.

To understand whether the RIM-BP2 phenotypes described so far are species or synapse type dependent, we chose to examine RIM-BP2 function at mouse hippocampal mossy fiber (MF) synapses, a mammalian synapse with distinct physiologically and anatomically properties. Notably, MF synapses strongly facilitate but possess multiple release sites^14^.

Our recordings in acute hippocampal brain slices revealed that neurotransmission at MF synapses is severely impaired upon the loss of RIM-BP2, compared to that at CA3-CA1 synapses. Furthermore, we also show that RIM-BP2 loss leads to a defective recruitment of Munc13-1 to the active zone specifically at MF synapses, but not at CA3-CA1 synapses, indicative of diversified functions of RIM-BP2 at these two synapse types. While at CA3-CA1 synapses RIM-BP2 maintains high fidelity coupling of Ca^2+^-channels to release sites, at MF synapses RIM-BP2 is required for proper docking and priming of synaptic vesicles to release sites, a yet undescribed function. Finally, our analysis of the active zone architecture reveals that RIM-BP2 and Munc13-1 clusters as well as RIM1 and Cav2.1 clusters are positioned at increased distances in MF synapses compared to that in CA3-CA1 synapses, demonstrating that these synapses utilize different architectural organizational principles.

## Results

### Distinct role of RIM-BP2 at hippocampal synapses

To probe the nature of diversity between central mammalian synapses, we examined the role of RIM-BP2 throughout the hippocampus. Immunostainings for RIM-BP2 in mouse hippocampal slices revealed RIM-BP2 expression in the whole hippocampal neuropil, with a strong labeling of the mossy-fiber layer band in stratum lucidum of area CA3 (Fig. 1a).

**Figure 1:**
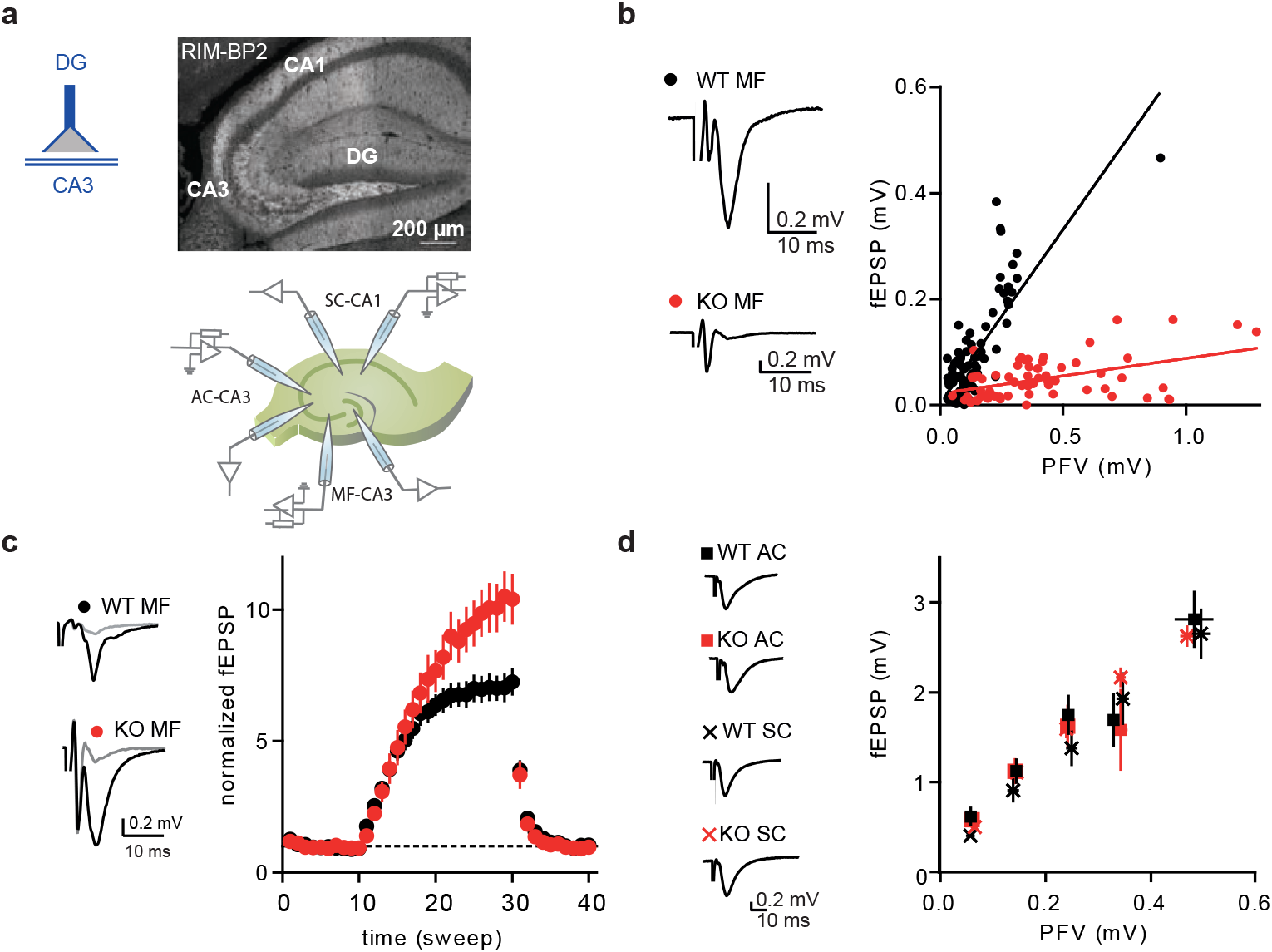
RIM-BP2 KO impacts synaptic transmission specifically at MF synapses. (**a**) Immunostaining of RIM-PB2 in hippocampal brain slices (DG = dentate gyrus) and schematic illustration of recording configurations. (**b**) Input-output of synaptic transmission of MF synapses, plotted as PFV against fEPSP amplitude. Sample traces show averages of 10 sweeps. (**c**) Frequency facilitation with 1 Hz stimulation of MF synapses (sweep 10 - 30). Sample traces show averages of five sweeps before (grey) and at the end of 1 Hz stimulation (black). (**d**) Input-output of synaptic transmission, plotted as PFV against fEPSP amplitude, of associative commissural (AC) and Schaffer collateral (SC) synapses showed no difference between RIM-BP2 WT and KO slices. Sample traces show averages of 10 sweeps. Values represent mean ± SEM.

To examine the functional impact of loss of RIM-BP2 at different hippocampal synapses, we recorded field excitatory postsynaptic potentials (fEPSPs) in acute brain slices obtained from RIM-BP2 KO mice. To ensure MF origin, we verified input sensitivity to group II metabotropic glutamate receptor (mGluR) agonist DCG IV^15^. The ratio of fEPSP to presynaptic fiber volley (PFV) was drastically reduced when stimulating the MF pathway in RIM-BP2 deficient (KO) slices compared to that in wildtype (WT) slices (Fig. 1b). This shows that neurotransmission is severely impaired at RIM-BP2 KO MF synapses. Additionally, we found that 1 Hz facilitation was significantly enhanced at MF synapses from RIM-BP2 deficient mice, suggesting a role for RIM-BP2 in short-term plasticity at the MF synapse (Fig. 1c). In contrast, the ratio of fEPSP to presynaptic fiber volley (PFV) for associative commissural (AC)-fibers and Schaffer collaterals (SC), both representing small central synapses, were not affected by loss of RIM-BP2 (Fig. 1d). Thus, RIM-BP2 deletion specifically impairs neurotransmitter release at hippocampal MF synapses, compared to AC and SC synapses.

### RIM-BP2 deletion does not alter Ca^2+^-channel localization at the MF synapse

Given that RIM-BP2 contributes to the coupling of Ca^2+^-channels and release apparatus in CA3-CA1 synapses^10,11^, we speculated that a mislocalization of Ca^2+^-channels in RIM-BP2 KO MF synapses might contribute to the severe phenotype in neurotransmission (Fig. 1). To test this hypothesis, we determined the position of clusters formed by the P/Q type Ca^2+^-channel subunit Ca_v_2.1 relative to clusters formed by the active zone protein RIM1, and the postsynaptic scaffold Homer1 in RIM-BP2 WT and KO MF synapses using super-resolution time-gated stimulated emission depletion (gSTED) *in situ*. However, RIM-BP2-deficient MF synapses exhibited no difference in the number of Ca^2+^-channel clusters, (Fig. 2a-c) or in the distance between Ca_v_2.1 and RIM1 as compared to WT (Fig. 2d). Therefore, the strong defect in neurotransmitter release in RIM-BP2 KO MF synapses is not due to a delocalization of Ca^2+^-channels.

**Figure 2:**
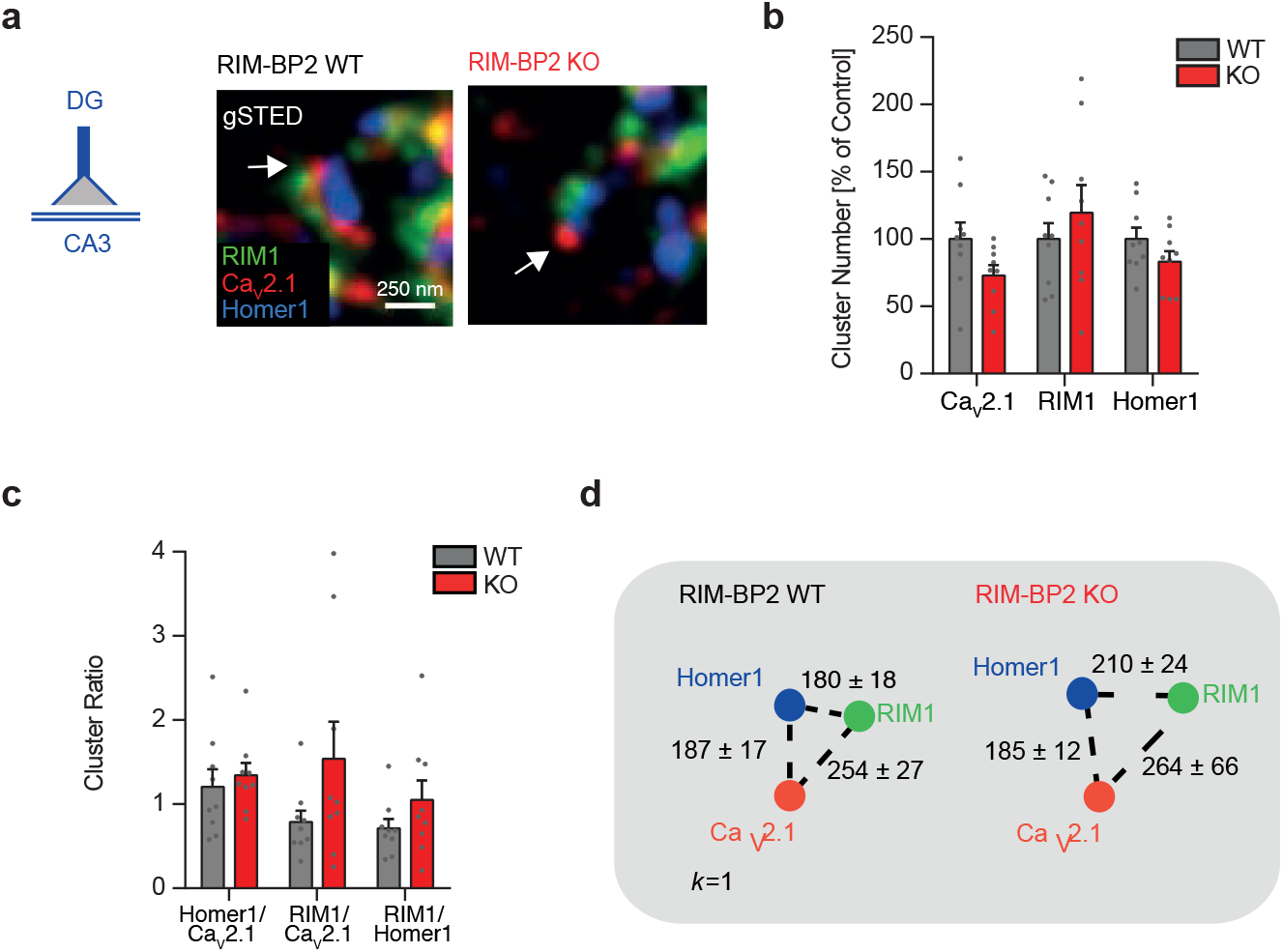
RIM-BP2 deletion does not alter the localization of Ca_V_2.1 clusters relative to the active zone protein RIM1 and the postsynaptic marker Homer1. (**a**) gSTED images of Ca_V_2.1, RIM1 and Homer1 clusters at MF boutons (MFBs) of RIM-BP2 WT and KO brain slices. (**b**) Cluster number of Ca_V_2.1, RIM1 and Homer1 at MFBs of RIM-BP2 WT and KO mice, normalized to WT mice. (**c**) Cluster ratios and distances of the first closest *k* neighbor (*k*=1,**d**), no significant differences were observed between RIM-BP2 WT and KO mice. Values represent mean ± SEM.

### MF synapses and CA3-CA1 synapses display differences in their active zone protein architecture

Given that RIM-BP2 KO does not alter Ca^2+^-channel localization at MF synapses but strongly decreases neurotransmitter release, we hypothesized that RIM-BP2 deletion might alter the abundance or localization of other presynaptic scaffold proteins at MF synapses. To test this hypothesis we performed a more detailed analysis of the positioning of scaffold protein clusters at WT MF synapses using gSTED. Interestingly, at WT MF synapses, the distance between RIM1 clusters and Ca_v_2.1 clusters was more than 35% larger (254 ± 27 nm) (Fig. 2d) than what we previously found at WT CA3-CA1 synapses^11^. In separate experiments, we also determined the localization of RIM-BP2 clusters relative to clusters of the priming/docking factor Munc13-1 and the scaffold protein Bassoon. Again, at WT MF synapses Munc13-1 clusters were positioned at increased distances (174 ± 20 nm) from RIM-BP2 clusters (Fig. S1a, b) than what we previously observed at CA3-CA1 synapses (115 ± 5 nm)^11^. Our results support the idea that MF synapses display a distinct active zone organization compared to CA3-CA1 synapses. These findings indicate that RIM-BP2 differentially impacts on neurotransmission at these two synapse types, possibly resulting from differences in their active zone protein architecture.

### RIM-BP2 docks synaptic vesicles *via* the specific recruitment of Munc13-1 at MF synapses

To investigate the possible mechanisms leading to the severe phenotype observed in RIM-BP2 deficient MF synapses, we compared the relative localization and abundance of the priming factor Munc13-1 and Ca_V_2.1 clusters in RIM-BP2 WT and KO brain slices by using gSTED. We found a drastic reduction in Munc13-1 cluster number (Fig. 3a-d), accompanied with an increased distance between Munc13-1 and Ca_v_2.1 clusters in RIM-BP2 deficient MF synapses compared to WT MF synapses (Fig. 3e,f), parameters which were unaltered in CA3-CA1 synapses (Fig. 3g-l). We also analyzed the relative distribution and abundance of Munc13-2 relative to Cav2.1 clusters at both MF and CA3-CA1 synapses. In both synapses, loss of RIM-BP2 neither altered the levels nor the distribution of Munc13-2 relative to Cav2.1 channels (Fig. S2). Therefore, RIM-BP2 is needed specifically in MF synapses to stabilize Munc13-1 at the active zone.

**Figure 3:**
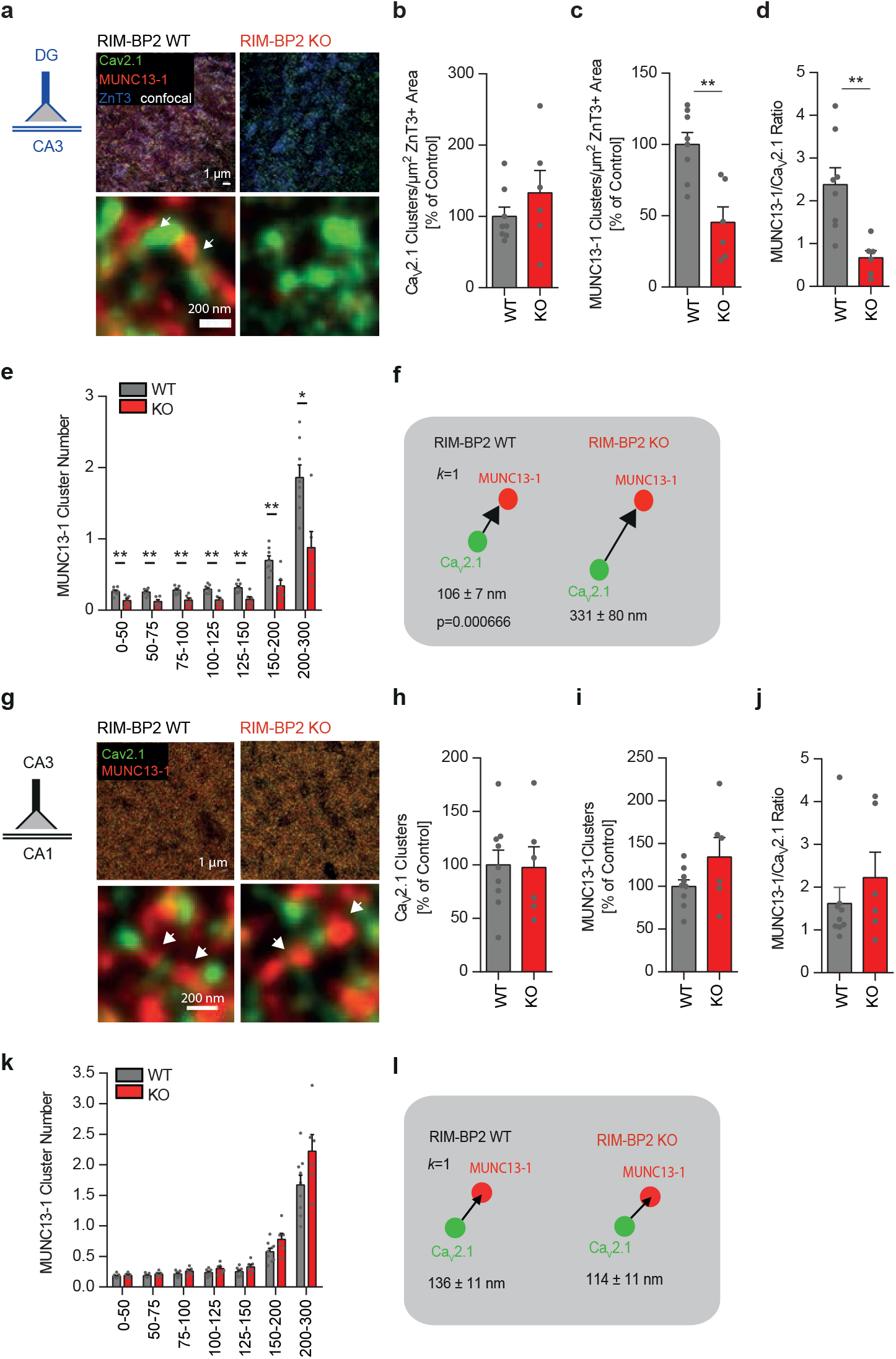
Loss of RIM-BP2 specifically reduces MUNC13-1 levels at MF synapses but not at CA3-CA1 synapses. (**a**) Representative gSTED images of Ca_V_2.1 and Munc13-1 clusters at MF boutons (MFB) identified by ZnT3 expression (confocal) in RIM-BP2 WT and KO brain sections. Arrows indicate Munc13-1 clusters nearby Ca_V_2.1 clusters. (**b**) Number of Ca_V_2.1 clusters per µm^2^ of ZnT3+ area in RIM-BP2 KO and WT mice, normalized to RIM-BP2 WT. (**c**) Number of Munc13-1 clusters per µm^2^ of ZnT3+ area normalized to RIM-BP2 WT mice. (**d**) Ratio of MUNC13-1 clusters/Ca_V_2.1 clusters in RIM-BP2 KO and WT mice. (**e**) The number of Munc13-1 clusters at determined distance intervals (nm) from a given Ca_V_2.1 cluster decreased significantly at all distances analyzed in RIM-BP2 KO, while the distance of the first closest *k* neighbor (*k*=1; **f**) significantly increased. (**g**) Representative gSTED images of Ca_V_2.1 and Munc13-1 clusters at CA3-CA1 synapses in RIM-BP2 WT and KO brain sections. Arrows indicate Munc13-1 clusters nearby Ca_V_2.1 clusters. (**h**) Number of Ca_V_2.1 clusters and MUNC13-1 clusters (**i**) found at CA3-CA1 synapses in RIM-BP2 KO and WT mice, normalized to RIM-BP2 WT mice. (**j**) Ratio of MUNC13-1 clusters/Ca_V_2.1 clusters at CA3-CA1 synapses. (**k**) At CA3-CA1 synapses, loss of RIM-BP2 does not significantly alter either the number of Munc13-1 clusters at determined distance intervals (nm) from a given Ca_V_2.1 cluster or the distance at which the first closest *k* neighbor (*k*=1, **l**) is found. Values represent mean ± SEM. *p<0.05, **p<0.01.

The major role of Munc13-1 is to dock and prime vesicles at the active zone^16,17^. To explore whether the loss of Munc13-1 at RIM-BP2 deficient MF synapse affected vesicle docking, we performed high-pressure cryo-fixation of acute hippocampal slices from WT and RIM-BP2 deficient animals and analyzed MF active zone structures by using transmission electron microscopy (EM). Indeed, EM images of RIM-BP2 deficient MF synapses exhibited a nearly 50% reduction in docked vesicles and a reduction in vesicle number within 30 nm from the active zone membrane (Fig. 4a,b), whereas the number of docked vesicles in CA1 synapses was unaltered (Fig. 4c,d). Thus, in contrast to CA3-CA1 synapses, MF synapses require RIM-BP2 dependent stabilization of Munc13-1 at the active zone to dock synaptic vesicles.

**Figure 4:**
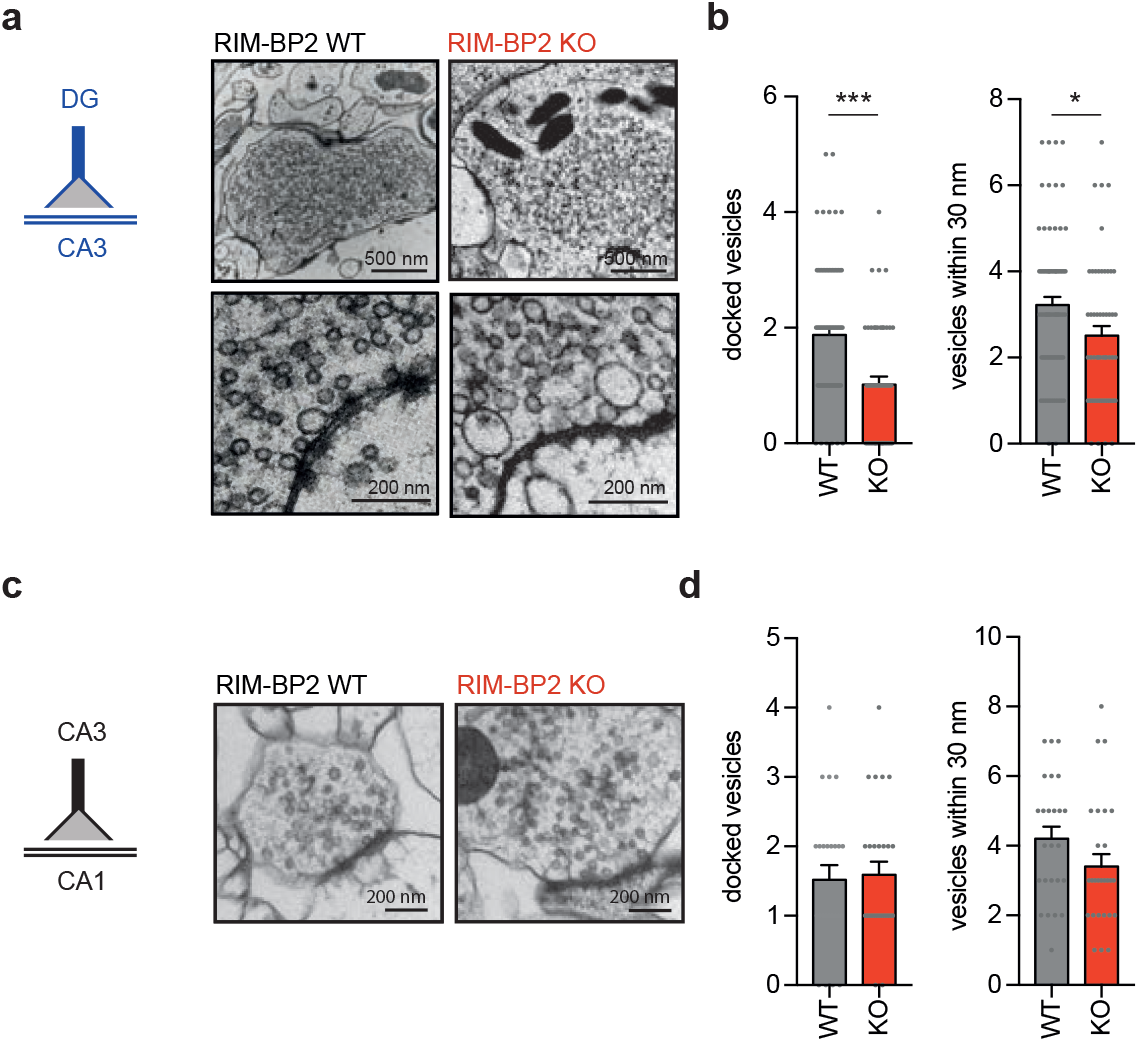
Loss of RIM-BP2 specifically impacts vesicle docking at MF synapses. (**a**) Representative EM images of MF synapses from acute hippocampal slices obtained from RIM-PB2 KO and WT mice. (**b**) Summary graphs show a reduction of docked vesicles and vesicles within 30 nm of the active zone at RIM-BP2 KO MF synapse compared to WT MF synapses. (**c**) Representative EM images of CA3-CA1 synapses of acute hippocampal slices from RIM-BP2 KO and WT mice. (**d**) Summary graph of docked vesicles and vesicles within 30 nm of the active zone show no difference for RIM-BP2 KO and WT synapses. Values represent mean ± SEM. *p<0.05, **p<0.01.

### Loss of RIM-BP2 impairs vesicle priming and release efficiency in granule autaptic neurons

To identify the molecular mechanism of RIM-BP2 function in MF synapses, we prepared autaptic cultures of hippocampal granule cells that form synaptic contacts only with themselves and therefore allow the quantitative analysis of synaptic input-output properties. It is important to note that cultured granule cells, like those in acute hippocampal brain slice, form MF like boutons as assessed by EM analysis (Fig. S3a). Additionally, autaptic granule cells are sensitive to DCG IV and thus can be pharmacologically identified (Fig. 5a,b S3). This shows that many aspects of the hippocampal granule cell identity, including the gross MF synapse structure and presynaptic mGLUR2 expression, are likely to be intrinsically encoded and independent of post-synaptic targets.

**Figure 5:**
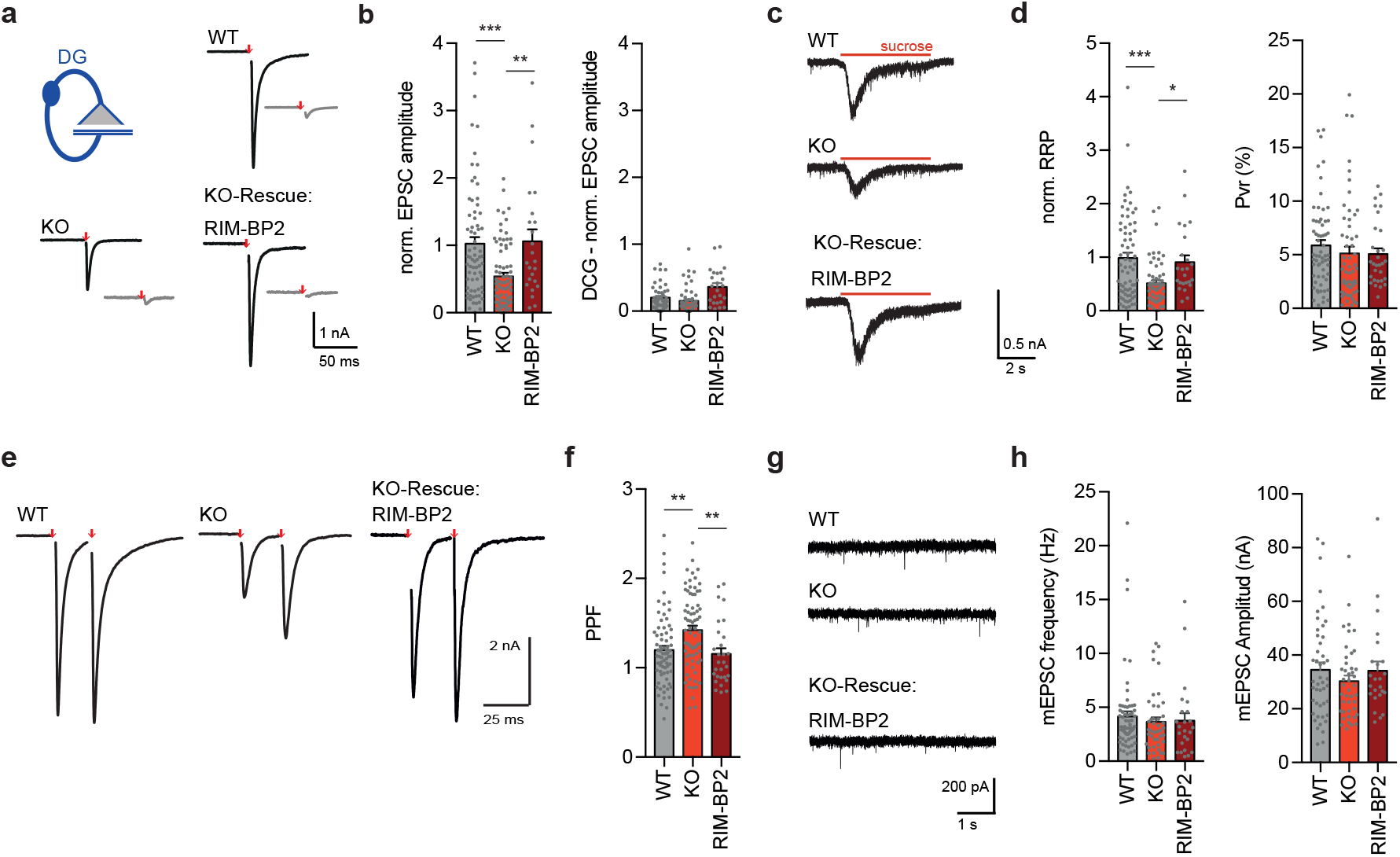
RIM-BP2 KO impacts synaptic transmission at granule autaptic neurons. (**a**) Sample traces of evoked EPSC amplitudes before (black) and after DCG IV application (grey) for RIM-BP2 WT and KO neurons. RIM-BP2 KO neurons were rescued by lentiviral transduction of RIM-BP2. (**b**) Summary graphs of normalized EPSC amplitudes evoked by 2 ms depolarization (red arrow). (**c**) Sample traces and (**d**) summary graphs of normalized RRP responses elicited by a 5 s application of 500 mM sucrose. Summary graph of the P_VR_ calculated as the ratio of the EPSC charge and the RRP charge. (**e**) Sample traces of evoked EPSC amplitudes with an interstimulus interval of 25 ms. (**f**) Summary graph of paired-pulse ratio (PPR) of RIM-BP2 WT, KO and RIM-BP2 rescued autaptic granule neurons. (**g**) Sample traces of miniature EPSCs (mEPSCs) and (**h**) summary graph of mEPSC amplitudes and frequencies. Values represent mean ± SEM. *p<0.05, **p<0.01, ***p<0.001.

Next, we wanted to test whether autaptic granule neurons show the same level of reduction in neurotransmitter release upon RIM-BP2 loss and thus whether they can be used as a model system for studying differences between the active zone protein architectures of diverse synapse types. Consistent with the findings from hippocampal field recordings (Fig. 1), evoked excitatory postsynaptic currents (EPSCs) were severely impaired in RIM-BP2 KO granule cell autapses compared to that in WT autapses (Fig. 5a,b). Rescue of RIM-BP2 deficiency by lentiviral expression in RIM-BP2 KO neurons completely restored synaptic transmission, confirming the specificity of RIM-BP2 function at these synapses (Fig. 5a,b). To examine the origin of the impaired evoked response, we first probed vesicle priming by measuring the readily releasable pool (RRP) *via* hypertonic sucrose solution application. In line with the finding that RIM-BP2 KO MF synapses had only half the number of docked vesicles compared to that in WT neurons (Fig. 4), the size of the RRP was reduced by 50 % as well (Fig. 5c,d). The frequency and amplitude of Ca^2+^-independent release events as measured by recording spontaneous miniature EPSCs (mEPSCs) was not significantly altered upon RIM-BP2 loss (Fig. 5g,h). Next, we investigated the paired-pulse ratio (PPR), a sensitive indicator of changes in release probability, and found that RIM-BP2 KO neurons displayed slightly enhanced facilitation compared to WT neurons (Fig. 5e,f). Furthermore, EPSCs evoked by 10 Hz action potential trains displayed less depression over the 5s train duration in RIM-BP2 KO granule cell autapses compared to WT or rescue groups (Fig. S4). Thus, using an additional model system, we could validate our initial findings and demonstrate that loss of RIM-BP2 leads to impaired hippocampal granule cell output, due to changes in both vesicle priming and release efficacy.

### RIM-BP2 primes synaptic vesicles *via* Munc13-1 in granule autaptic neurons

Our gSTED imaging experiments revealed a decrease in synaptic Munc13-1 localization upon the loss of RIM-BP2 (Fig. 3). In small central synapses, priming is attained by an interaction of Munc13 and RIM *via* the Munc13 C_2_A domain, which can be mimicked by the constitutively monomeric mutant Munc13-K32E lacking Munc13-1 homodimerization^18^. We therefore examined whether the priming deficit observed in RIM-BP2-deficient MF synapses could be rescued by restoring Munc13 function by expression Munc13-1 WT (M13^WT^) or the constitutively monomeric Munc13-1 mutant (M13^K32E^)^19^. Remarkably, Munc13-K32E expression in RIM-BP2 KO granule neurons sufficed to rescue vesicle priming and Ca^2+^-evoked neurotransmission (Fig. 6), whereas Munc13-1 WT was not sufficient to rescue the RIM-BP2 KO phenotype (Fig.6). This suggests that in MF synapses, unlike hippocampal pyramidal neuron synapses, the recruitment of active monomeric Munc13 is RIM-BP2 dependent.

**Figure 6:**
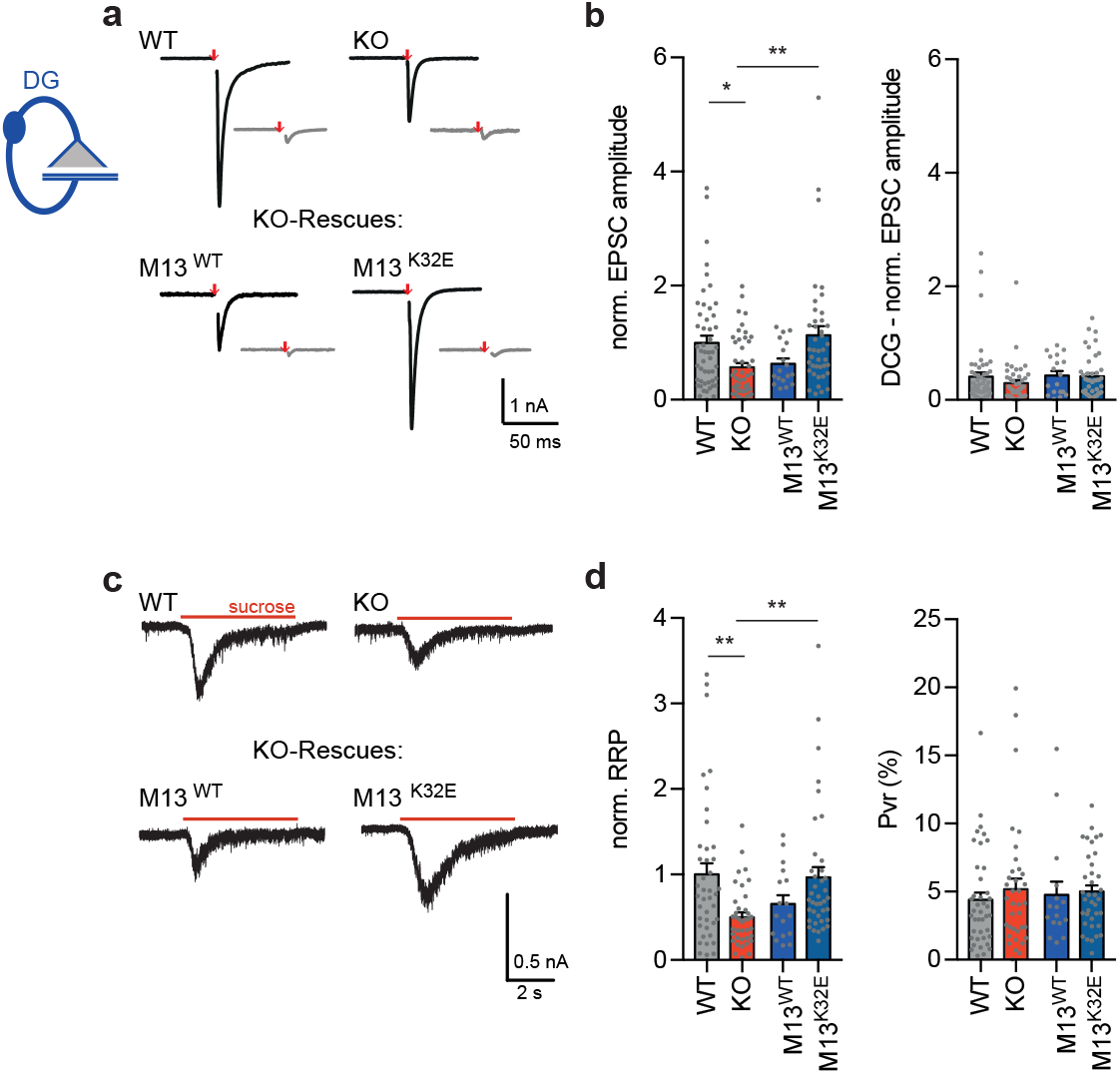
Monomeric Munc13-1 rescues vesicle priming in RIM-BP2 KO granule autaptic neurons. (**a**) Sample traces of evoked EPSC amplitudes before (black) and after DCG IV application (grey) for RIM-BP2 WT and KO neurons and lentiviral-transduced RIM-BP2 KO rescues with either Munc13-1 WT (M13^WT^) or Munc13-1 K32E (M13^K32E^). (**b**) Summary graphs of normalized EPSC amplitudes evoked by 2 ms depolarization (red arrow). (**c**) Sample traces and (**d**) summary graphs of normalized RRP responses elicited by a 5 s application of 500 mM sucrose. Summary graph of the P_VR_ calculated as the ratio of the EPSC charge and the RRP charge. Values represent mean ± SEM. *p<0.05, **p<0.01.

## Discussion

Chemical synapses have been highly diversified by evolution in their molecular composition, ultrastructure, and consequently function. In the last decades, presynaptic diversity was studied in synapses that exhibit ultrastructural differences, such as the T-bar structure in the neuromuscular junction of drosophila melanogaster or the ribbon synapse of vertebrate photoreceptor cells^13^. Ultrastructural diversity is often associated with the expression of specialized synaptic organizers, like Bruchpilot or RIBEYE, which shape presynaptic structure and function^13,20,21^. More recently, the heterogeneity of distinct synapses came into focus, since many central synapses seemingly express similar presynaptic proteins but still show distinct release probabilities and Ca^2+^-secretion coupling. Synaptic heterogeneity might be achieved by several mechanisms, including variation in the abundance of single proteins or expression of different protein isoforms. Notably, a contribution of the exact nanoscale arrangement of proteins at active zones to synaptic diversity has also been discussed^2,22^.

To understand the function of a given protein in neurotransmitter release, one can compile the results from protein knockouts in diverse synapses and extract a universal function. Some highly preserved proteins, like Munc13-1, are essential for neurotransmitter release in a variety of synapses, since they exclusively conduct one specific function, in this case vesicle priming^17,23–25^. However, many other presynaptic proteins show distinct knockout phenotypes for diverse synapses, like RIM-BPs. In small central synapses, the calyx of Held, and inner ear hair cells, RIM-BP2 deletion has rather minor effects on neurotransmitter release, which results only from a mild loosening of the coupling between Ca^2+^-channels and synaptic vesicles^10,11,26^. Strikingly, we now show that in large MF synapses, RIM-BP2 deletion strongly impairs neurotransmitter release due to an essential role of RIM-BP2 in recruiting Munc13-1 to the active zone and thereby promoting vesicle docking and priming. Therefore, RIM-BP2 function seems as diverse as the synapse type where it is expressed.

Why does RIM-BP2 function differ between hippocampal CA3-CA1 and MF synapses? Due to known differences in their anatomy and nanodomain coupling of Ca^2+^-channel^14,27^, one might speculate that different nanoscale arrangements in the active zone of CA3-CA1 and MF synapses might contribute to synaptic diversity. Indeed, here we show that under normal conditions the distances between the presynaptic scaffold protein clusters of RIM1 and Cav2.1, as well as between RIM-BP2 clusters and Munc13-1 clusters are increased in MF synapses compared to CA3-CA1 synapses.

Loss of RIM-BP in the drosophila NMJ displays a very severe phenotype, including impairment of synaptic vesicle docking and Ca^2+^-channel mislocalization^9^. In the mammalian hippocampus, these distinct functions of RIM-BP2 are differentially partitioned at two synapse types. Whereas RIM-BP2 deletion in CA3-CA1 synapses alters short-term synaptic plasticity by mild alterations in Ca^2+^-channel localization^11^, loss of RIM-BP2 at MF synapses reduced vesicle docking and priming due to a severe reduction in synaptic Munc13-1 protein levels. Importantly, in RIM-BP2 KO, we detected no significant change in Ca^2+^-channel localization at MF synapses, whereas Munc13-1 abundance and vesicle docking were unaffected in CA3-CA1 synapses. These results suggest that the protein structure of RIM-BP2 intrinsically encodes two different functions: Ca^2+^-channel localization and vesicle docking/priming *via* Munc13-1, two functions emerging in distinct active zone architectures. Therefore, the protein’s synaptic context seems important in determining the exact protein function. It will be of interest to see whether and through which molecular mechanism loss of RIM-BP2 alters the function of other mammalian synapses as well.

In mammalian synapses and drosophila NMJ, RIM-BPs have been shown to interact with Ca^2+^-channels^9,26^. While we previously observed mislocalization of Cav2.1 clusters within the AZ of CA3-CA1 synapses upon the loss of RIM-BP2, at MF synapses the closest distance at which Cav2.1 clusters are found relative to RIM1 and Homer1 clusters was unchanged. This possibly indicates that RIM-BP2 is not required to fine-tune Ca^2+^-channel localization at MF synapses. Interestingly, CA3-CA1 synapses are tight-coupled synapses^28^, whereas MF synapse depict loose Ca^2+^-channel release site coupling^27^, suggesting that RIM-BP2 might play different roles in Ca^2+^-secretion coupling at these two synapse types.

In CA3-CA1 synapses vesicle priming is accomplished by the interaction of RIM and Munc13-1^18,29^, which is not affected by the deletion of RIM-BP2^11^. However at MF synapses, RIM-BP2 deletion results in a severe reduction in synaptic Munc13-1 but not Munc13-2 protein levels and likely, as a direct consequence, provokes a deficit in vesicle docking and priming. These results indicate that RIM-BP2 promotes vesicle priming at MF synapses specifically *via* Munc13-1. The RIM-BP2 phenotype, which we describe here is similar to the knockout of Munc13-2 at MF synapses^30^. Nevertheless, in RIM-BP2 KO MF synapses, Munc13-2 levels as well as its localization to the Ca^2+^-channels are seemingly unaltered, indicating that at MF synapses RIM-BP2 is specifically required for Munc13-1 dependent vesicle priming.

One explanation for why RIM-BP2 is crucial for neurotransmitter release at MF synapses could be that due to increased distances between the scaffold proteins at MF terminals, the MF active zone requires additional protein interactions to sufficiently prime synaptic vesicles. RIM1 is known to interact with Munc13-1 and RIM-BP2, while at the same time RIM-BP2 has been reported to directly interact with RIM1 and Bassoon^26,31,32^. Additionally, we previously have shown that Munc13-1 co-immunoprecipitates with RIM-BP2 from synaptosome preparation, suggesting that these proteins act in the same protein complex^11^. Therefore, RIM-BP2 promotes Munc13-1 localization *via* RIM. Interestingly, RIM-BP2 function is obviously still dependent on RIM activity, since the overexpression of Munc13-1 K32E^18^, the constitutively active form of Munc13-1, but not Munc13-1 WT rescued the RIM-BP2 KO phenotype at MF synapses. Hence, the loss of RIM-BP2 at MF synapses reveals a hierarchical interaction between RIM-BP2, RIM1 and MUNC13-1, where RIM-BP2 recruits RIM1, which in turn monomerizes Munc13-1 to build the priming complex for synaptic vesicles. Our data suggest that RIM-BP2 increases the affinity for RIM to Munc13-1 and consequently stabilizes Munc13-1 at presynaptic active zones. Notably, RNA-seq analysis points towards lower RIM1 levels at MF synapses than at CA3-CA1 synapses^33^. Therefore, stabilization of RIM1 *via* RIM-BP2 might be more important at MF synapses.

We see two possibilities for why RIM-BP2 function is redundant for CA3-CA1 synapse neurotransmitter release: On the one hand, RIM-BP2 might be substituted by another protein and therefore possess a molecular redundancy. On the other hand, it might be structurally redundant, if the synaptic zone architecture itself at CA3-CA1 synapses enhances the affinity of RIM1 and Munc13-1 as a priming complex and is therefore RIM-BP2 independent. In any case, the functional hierarchy of the triple complex formed by RIM-BP2/RIM1/Munc13-1 is fundamentally different between CA3-CA1 and MF synapses. At MF synapses RIM-BP2 acts first to stabilize RIM1/Munc13-1 at the active zone, whereas at CA3-CA1 synapses, RIM-BP2 impacts synaptic function after RIM/Munc13-1 primes vesicles at the active zone.

Whereas RIM-BP function is essential for both Ca^2+^-secretion coupling and docking at the drosophila NMJ^9^, it seems that at hippocampal synapses, RIM-BP2 has diversified functions that seem to depend on the exact composition of the respective active zone. So far, factors determining synaptic diversity remain largely unknown. Importantly, we could show that MF synapses form in autaptic culture and maintain their functional properties. This suggests that yet unknown active zone super-organizers, such as scaffold proteins, intrinsically encode the diversity of synapse function, independently from the post-synaptic partner. Therefore, we propose that proteins, which are not absolutely essential for vesicular release in small central synapses, for example Liprins or SYD-1, may contribute to synaptic diversity and should be of particular interest for future studies.

## Material and Methods

### KO Mouse Generation

RIM-BP2 KO mouse generation and genotyping was performed as described previously^11^. All animal experiments were approved by the animal welfare committee of Charité Universitaetsmedizin Berlin and the Landesamt für Gesundheit und Soziales Berlin and carried out under the license (Berlin State Government, T0410/12; T0100/03).

### Slice Preparation and Electrophysiological Recordings

Acute hippocampal slices were prepared as described previously^11^. In brief, RIM-BP2 KO mice and wild-type littermates of both sexes (4-8 weeks) were anesthetized with Isofluorane and decapitated. The brain was quickly removed and chilled in ice-cold sucrose-artificial cerebrospinal fluid (sACSF) containing (in mM): 50 NaCl, 25 NaHCO3, 10 glucose, 150 sucrose, 2.5 KCl, 1 NaH2PO4, 0.5 CaCl2, and 7 MgCl2. All solutions were saturated with 95% (vol/vol) O2/5% (vol/vol) CO2, pH 7.4.

Slices (300 μm, sagittal or horizontal) were cut with a Leica VT1200S microtome (Wetzlar, Germany) and stored submerged in sACSF for 30 min at 35 °C and subsequently stored in ACSF containing (in mM): 119 NaCl, 26 NaHCO3, 10 glucose, 2.5 KCl, 1 NaH2PO4, 2.5 CaCl2 and 1.3 MgCl2 saturated with 95% (vol/vol) O2/5% (vol/vol) CO2, pH 7.4, at RT. Experiments were started 1 to 6 h after the preparation.

Experiments were conducted in parallel on a comparable number of slices from WT and KO animals prepared at the same experimental day for at least 3 times (biological replicates). Technical replicates were obtained for evoked responses and averaged.

For recordings, slices were placed in a recording chamber continuously superfused with ACSF at RT at a rate of 2.5 ml/min. fEPSPs were evoked by electrical stimulation with patch pipettes filled with ACSF. fEPSPs were recorded with a low-resistance patch-pipette filled with ACSF. Recordings were performed with a MultiClamp 700B amplifier. Signals were filtered at 2 kHz and digitized (BNC-2090; National Instruments Germany GmbH) at 10-20 kHz. IGOR Pro software was used for signal acquisition (WaveMetrics, Inc.).

For Mossy fiber recordings, stimulation electrodes were placed in the granule cell layer or in the hilus region. Mossy fiber origin of recorded signals was verified by frequency facilitation >400% when stimulus frequency was changed from 0.05 to 1 Hz and a complete block of responses upon DCG IV (1 µM; Tocris) application at the end of each experiment. fEPSPs in area CA1 were recorded in stratum radiatum after stimulation of the Schaffer collaterals. fEPSPs of associative commissural fibers in area CA3 were recorded in stratum radiatum after stimulation electrodes were placed in stratum radiatum, in the presence of DCG IV (1 µM) to avoid mossy fiber contamination. fEPSP magnitude was determined by analyzing ± 2 ms of the amplitude peak. Data were analyzed with the Igor plug-in NeuroMatic (neuromatic.thinkrandom.com) software. Recordings were only analyzed if the fiber volley remained constant throughout the recording. Statistical analysis was performed with Prism 6 (GraphPad Software).

### Autaptic Granule Cell Culture

Autaptic cultures of Dentate Gyrus Granule cells were prepared as described previously^34^. In brief, the dentate gyrus of P0-P1 RIM-BP2 WT and KO embryos was separated from the hippocampus. After digestion with Papain and trituration, cells were plated on astrocytic micro-islands^35^. Neurons were incubated at 37°C for 14-20 days before the electrophysiological characterization was performed. For rescue experiments, neurons were transduced with lentiviruses 24 hours after plating.

### Lentiviral Constructs

Lentiviral constructs used in this study were based on the FUGW vector^36^. The cDNA from mouse RIM-BP2 (NM_001081388) and from rat *Unc-13a* (NM_022861)^19^ were cloned into an lentiviral shuttle vector after a NLS-GFP-P2A or NLS-GFP-P2A under the control of a human *synapsin-1* promoter. The expression of nuclear RFP or GFP allows to identify transduced neurons. All lentiviruses were provided from the Viral Core Facility of the Charité Berlin and prepared as described before^36^.

### Electrophysiological Recordings of Autaptic Neurons

To pharmacologically identify autaptic granule cells, DCG IV (1µm) was washed in after each experiment. Only cells where synaptic transmission was inhibited by approximately 70 % were considered for analysis^34^.

Whole-cell patch-clamp recordings in autaptic neurons were performed as described previously^11^ at 13-21 days *in vitro* (DIV) with a Multiclamp 700B amplifier (Molecular Devices). Data were acquired from at least 3 different cultures (biological replicates). To minimize variability in recordings, about the same number of autapses were recorded from each experimental group each day. Technical replicates were obtained for evoked responses and averaged. Data were normalized to the mean value of the control group of each culture.

The patch pipette solution contained the following (in mM): 136 KCl, 17.8 HEPES, 1 EGTA, 4.6 MgCl_2_, 4 Na_2_ATP, 0.3 Na_2_GTP, and 12 creatine phosphate, and 50 U/ml phosphocreatine kinase (300 mOsm; pH 7.4). The recording chamber was constantly perfused with extracellular solution containing 140 mM NaCl, 2.4 mM KCl, 10 mM Hepes, 2 mM CaCl_2_, 4 mM MgCl_2_, and 10 mM glucose (pH adjusted to 7.3 with NaOH, 300 mOsm). Solutions were applied using a fast-flow system. Data were filtered at 3 kHz, digitized at 10 kHz, and recorded with pClamp 10 (Molecular Devices). Data were analyzed offline with Axograph X (AxoGraph Scientific) and Prism 6.

EPSCs were evoked by a 2-ms depolarization to 0 mV from a holding potential of −70 mV. PPRs were calculated as the ratio from the second and first EPSC amplitudes with an interstimulus interval of 25 ms. The RRP size was calculated by integrating the transient current component of 5 s evoked by application of extracellular hypertonic 500 mM sucrose solution. Miniature EPSC (mEPSC) amplitude and frequency were detected using a template-based algorithm in Axograph X.

### Immunohistochemistry, time gated STED microscopy and cluster distance analysis

Immunohistochemistry was performed as described previously (Grauel et al., 2016). Conventional confocal tile scans of RIM-BP2 immunofluorescence in the hippocampus were acquired with a Leica SP8 laser confocal microscope equipped with a 20× 0.7-N.A. oil immersion objective.

Following immunostaining, sagittal cryosections (10 μm) of RIM-BP2 WT and KO brains were imaged by gSTED with a Leica SP8 gSTED microscope (Leica Microsystems) as described previously (Grauel et al., 2016). Within each independent experiment, RIM-BP2 KO and WT samples were imaged with equal settings. Raw dual- and triple-channel gSTED images were deconvolved with Huygens Professional software (Scientific Volume Imaging) using a theoretical point spread function automatically computed based on pulsed- or continuous-wave STED optimized function and the specific microscope parameters. Default deconvolution settings were applied.

Experiments were repeated at least two times on different mice (biological replicates).

For cluster distance analysis, deconvolved images were threshold and segmented by watershed transform with Amira software (Visualization Sciences Group) to identify individual clusters and to obtain their x and y coordinates. Within the same independent experiment, the same threshold and segmentation parameters were used. According to the lateral resolution achieved, clusters with a size smaller than 0.0025 μm^2^ were not considered for analysis. To select MUNC13-1 and Ca_V_2.1 clusters within the ZnT3+ area, a mask was created applying a threshold on deconvolved ZnT3+ confocal images with Amira software (Visualization Sciences Group). The average number of clusters at specific distances and the k-nearest neighbor distance were analyzed with a MATLAB custom-written script, as previously described (Grauel et al., 2016).

### Electron Microscopy

Acute Hippocampal slices (150 µm) were prepared as described above and frozen at RT using an HPM 100 (Leica) supported with extracellular solution containing 15% Ficoll.

Slices from at least 3 different WT and KO animals were frozen and processed in parallel (biological replicate). After freezing, samples were transferred into cryovials containing 1% glutaraldehyde, 2% osmium tetroxide, and 1% ddH_2_O in anhydrous acetone in an AFS2 (Leica) with the following temperature program: −90°C for 72 h, heating to −60°C in 20 h, −60°C for 8 h, heating to −30°C in 15 h, −30°C for 8 h, heating to −20°C in 8 h. After staining with 1% uranyl acetate, samples were infiltrated and embedded into Epon and backed 48 h at 60 °C. Serial 40-nm sections were cut using a microtome (Leica) and collected on formvar-coated single-slot grids (Science Services GmbH). Before imaging, sections were contrasted with 2.5% (wt/vol) uranyl acetate and lead citrate. Samples were imaged in a FEI Tecnai G20 TEM operating at 80-120 keV and images taken with a Veleta 2K × K CCD camera (Olympus) and analyzed with a custom-written ImageJ (NIH) and MATLAB (The MathWorks, Inc.) script.

### Statistical Analysis

For electrophysiological experiments in brain slices, numbers of experiments are indicated in n/N, while n represents the number of brain slices and N the number of animals. Sample size estimation was done as published previously^30^.

For gSTED, statistical analysis was done with SPSS Statistics software (IBM). Normality was assessed checking histograms and Q-Q plots. Pairwise comparisons were analyzed with the Mann-Whitney *U* test. Significance threshold α was set to 0.05. Only p values less than 0.05 were considered significant. Values corresponding to one WT animal measured displaying an extreme outlier were excluded from the whole MF-CA3 MUNC13-1/Ca_V_2.1 data analysis, based on SPSS estimation of extreme values (value > Q3 + 3*IQR). Values are expressed as mean ± SEM, and *n* indicates the number of animal tested. Sample size estimation was done as published previously^11^.

For autaptic electrophysiological experiments, statistical analysis was done in Prism (Graphpad). First, the D’Agostino-Pearson test was applied to check whether data are normally distributed. If data were normally distributed, statistical significance was determined by using one-way ANOVA followed by Turkey *post hoc* test. For data which were not normally distributed, statistical significance was assessed using non-parametric Kruskal-Wallistest with Dunn’s *post hoc* test. Values are expressed as mean ± SEM, and *n* indicates the number of recorded autapses. Sample size estimation was done as published previously^*19,37*^.

For electron microscopy experiments, the D’Agostino-Pearson omnibus test was used to check for normal distribution of data. The For WT vs. KO comparison, an unpaired *t* test with Welch’s correction was used for normally distributed data and the Mann-Whitney *U* test was used for not normally distributed data. Values are expressed as mean ± SEM, and *n* indicates the number of active zones analyzed. Sample size estimation was done as published previously^11^.

## Acknowledgement

This work was supported by the Deutsche Forschungsgemeinschaft (Collaborative Research Grant SFB 958 [to D.S. (A5), S.S. (A3, A6), C.R. (A5)] and Excellence Cluster Neurocure Exc257 (to D.S., S.S., and C.R.). We thank Melissa Herman and Gülcin Vardar for providing critical advice for writing the manuscript and Alexander Walter for providing the gSTED analysis script and comments. We thank Berit Söhl-Kielczynski, Anke Schönherr, Susanne Rieckmann und Lisa Züchner for technical support; and Jörg Breustedt for discussions. We thank the Charité viral core facility for virus production and the cellular imaging facility of the Leibniz-Forschungsinstitut für Molekulare Pharmakologie (FMP) for the use of the gSTED microscope. We thank Anna Fejtova and Eckart Gundelfinger for the RIM-BP2 antibody. We thank Martin Lehmann (FMP Berlin, Cellular Imaging Facility) for the use of the Leica gSTED microscope.

## Author Contributions

M.M.B. designed and performed EM experiments and granule cell recordings

M.M. designed and performed gSTED experiments

C.W. designed and performed slice experiments M.M.B., C.R., M.M., S.S. and D.S. wrote the manuscript A.S., L.M.V. and P.B. assisted with slice recordings

B.B.A. and K.G. assisted with granule cell experiments

S.S., C.R. and D.S. designed experiments, supervised the project

---

**Figure S1:**
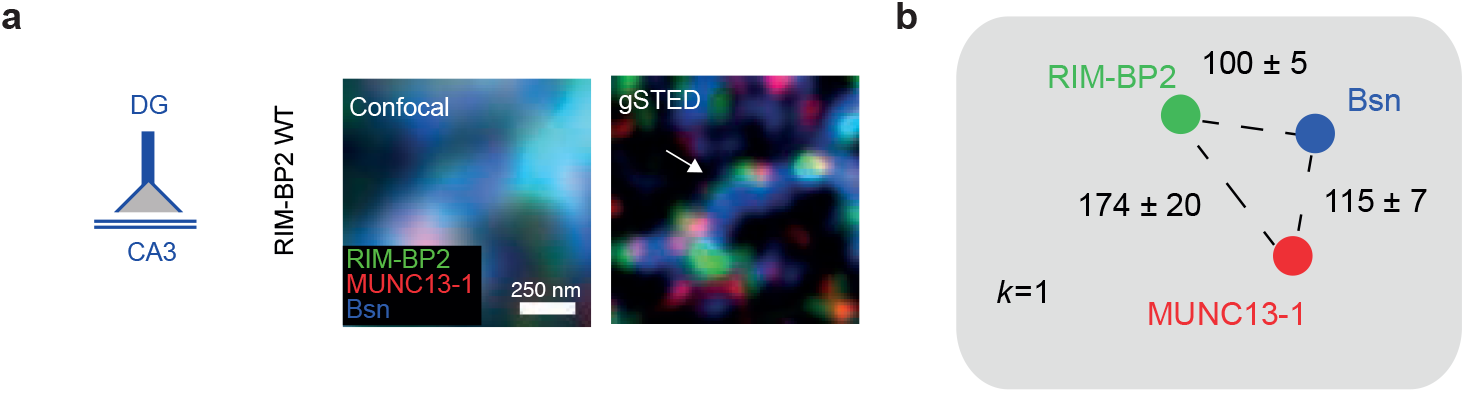
RIM-BP2 localization relative to MUNC13-1 and Bassoon at MF synapses. (**a**) Confocal (left) and gSTED (right) images of RIM-BP2, Munc13-1 and Bassoon (Bsn) at the active zone of WT MFBs *in situ*. (**b**) Distances at which the first closest *k* neighbor was found (nm). Values are mean ± SEM.

**Figure S2:**
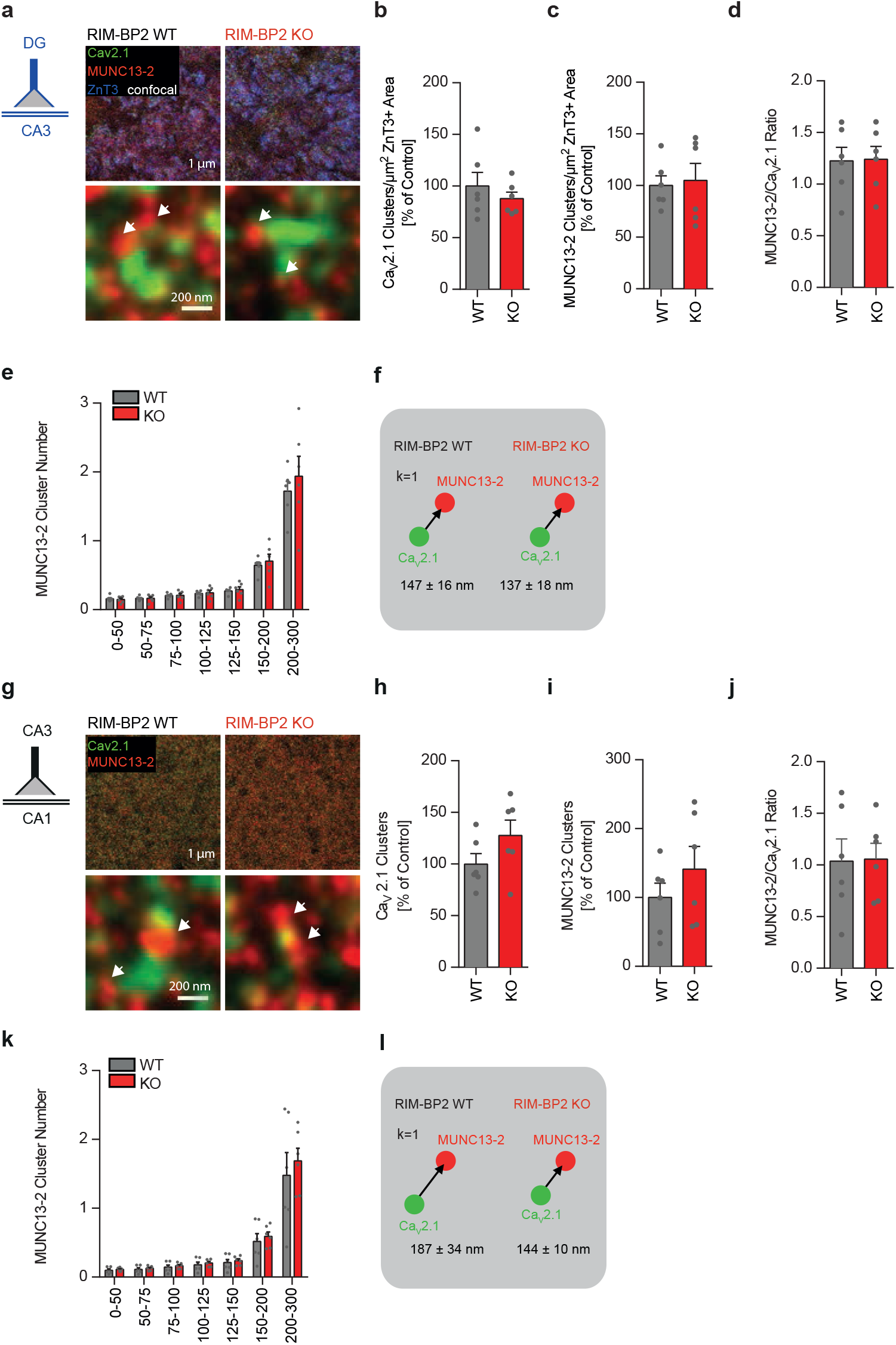
Loss of RIM-BP2 does not alter MUNC13-2 levels at both MF-CA3 and CA3-CA1 synapses. (**a**) Representative gSTED images of Ca_V_2.1 and Munc13-2 clusters at MF boutons (MFB) identified by ZnT3 expression (confocal) in RIM-BP2 WT and KO brain sections. Arrows indicate Munc13-2 clusters nearby Ca_V_2.1 clusters. (**b**) Number of Ca_V_2.1 clusters per µm^2^ of ZnT3+ area in RIM-BP2 KO and WT mice, normalized to RIM-BP2 WT mice. (**c**) Number of Munc13-2 clusters per µm^2^ of ZnT3+ area normalized to RIM-BP2 WT mice and ratio of MUNC13-2 clusters/Ca_V_2.1 clusters (**d**) in RIM-BP2 KO and WT mice. (**e**) Number of Munc13-2 clusters at determined distance intervals (nm) from a given Ca_V_2.1 cluster and distance at which the first closest *k* neighbor (*k*=1; **f**) is found. (**g**) Representative gSTED images of Ca_V_2.1 and Munc13-2 clusters at CA3-CA1 synapses in RIM-BP2 WT and KO brain sections. Arrows indicate Munc13-2 clusters nearby Ca_V_2.1 clusters. (**h**) Number of Ca_V_2.1 clusters and MUNC13-2 clusters (**i**) found at CA3-CA1 synapses in RIM-BP2 KO and WT mice, normalized to RIM-BP2 WT mice. (**j**) Ratio of MUNC13-2 clusters/Ca_V_2.1 clusters at CA3-CA1 synapses. (**k**) At CA3- CA1 synapses, loss of RIM-BP2 does not significantly alter either the number of Munc13-1 clusters at determined distance intervals (nm) from a given Ca_V_2.1 cluster or the distance at which the first closest *k* neighbor (*k*=1) is found.

**Figure S3:**
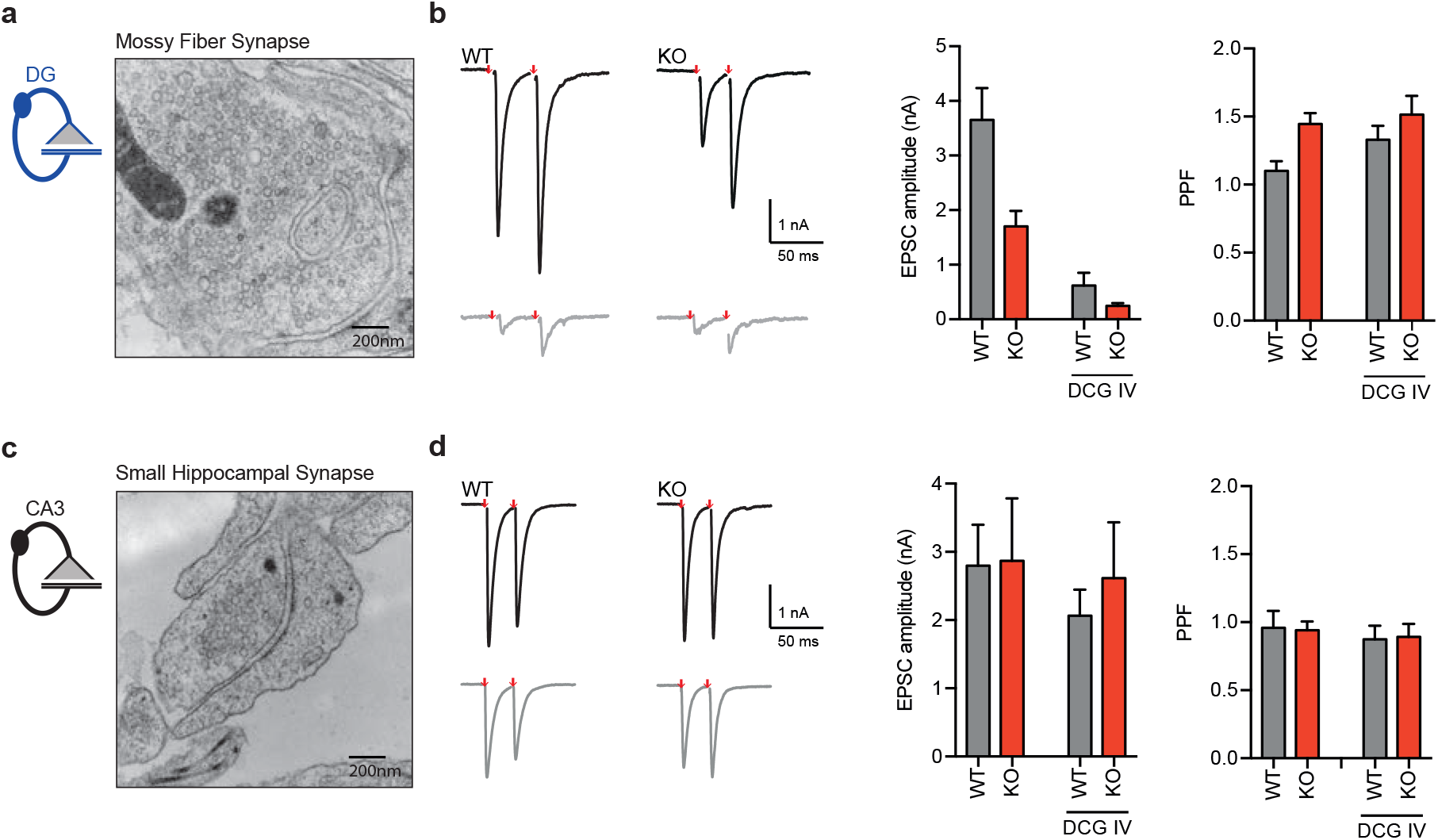
Characterization of autaptic granule cells. (**a**) Representative EM image of a granule cells in culture. (**b**) Sample traces and summary graphs of evoked EPSCs of granule cells with an interstimulus interval of 25 ms before (black) and after DCG IV application (grey) of RIM-BP2 WT and KO neurons. Both WT and KO neurons respond to the DCG IV application. EPSC amplitudes were reduced in RIM-BP2 KO neurons. (**c**) Representative EM image of a small central synapse in culture. (**d**) Sample traces and summary graphs of evoked EPSCs of hippocampal autaptic neurons with an interstimulus interval of 25 ms from RIM-BP2 WT and KO neurons that did not respond to DCG IV application. Autaptic neurons that do not respond to DCG IV application showed no reduction in EPSC amplitude in RIM-BP2 KO autapses compared to WT.

**Figure S4:**
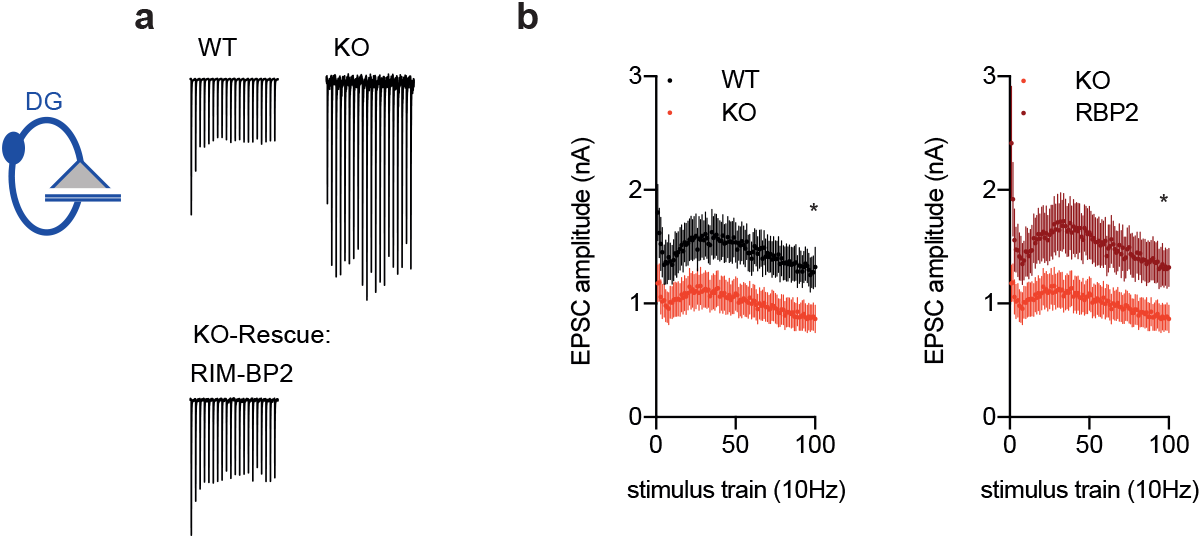
Loss of RIM-BP2 increases synaptic facilitation in autaptic granule neurons. **(a)** Sample traces and (**b**) summary graph of EPSCs elicited by a 10 Hz stimulation train normalized to the first EPSC. Values represent mean ± SEM. *p<0.05

**Table S1.** Primary and secondary antibodies for immunohistochemistry.

| **Antigen** | **Species** | **Dilution** | **Source** |
| --- | --- | --- | --- |
| Bsn<br>(N-terminal) | ms | 1:1200 (IHC) | Abcam |
| Cav2.1<br>(rat aa1921-2212) | Rb | 1:500 (IHC) | Synaptic System |
| Cav2.1<br>(rat aa1921-2212) | GP | 1:500 | Synaptic System |
| Homer1<br>(human aa 1-186) | GP | 1:200 (IHC) | Synaptic System |
| MUNC13-1<br>(rat aa 3-317) | Rb | 1:150 (IHC) | Synaptic System |
| MUNC13-2<br>(rat aa 151-317) | Rb | 1:150 (IHC) | Synaptic System |
| RIM1<br>(rat aa 602-723) | ms | 1:200 (IHC) | BD Pharmigen |
| RIM-BP2 (rat aa 589-869) | GP | 1:600/1:1000 (IHC) | Kind gift of A. Fejtova and<br>Eckart Gundelfinger<br>(Leibniz Institute for<br>Neurobiology, Magdeburg,<br>Germany) |
| ZnT3<br>(mouse aa 2-75) | ms | 1:500 | Synaptic System |
| Anti mouse AF488 | goat | 1:100/1:200<br>(gSTED)<br>1:400 (confocal) | Invitrogen |
| Anti guinea pig AF594 | goat | 1:100/1:200<br>(gSTED)<br>1:400 (confocal) | Invitrogen |
| Anti rabbit ATTO647N | goat | 1:100 (gSTED) | Active Motif |

**Table S2:**
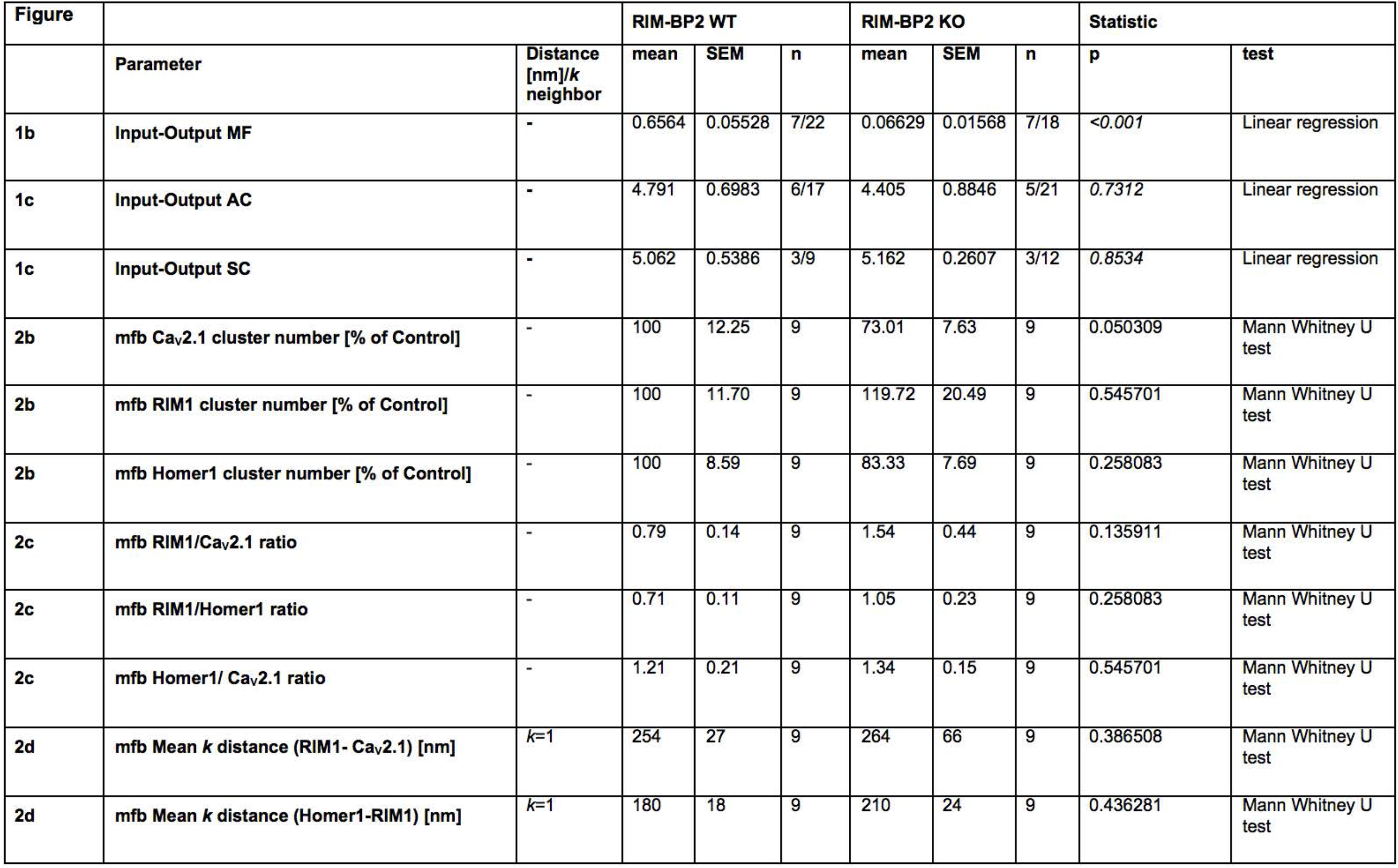
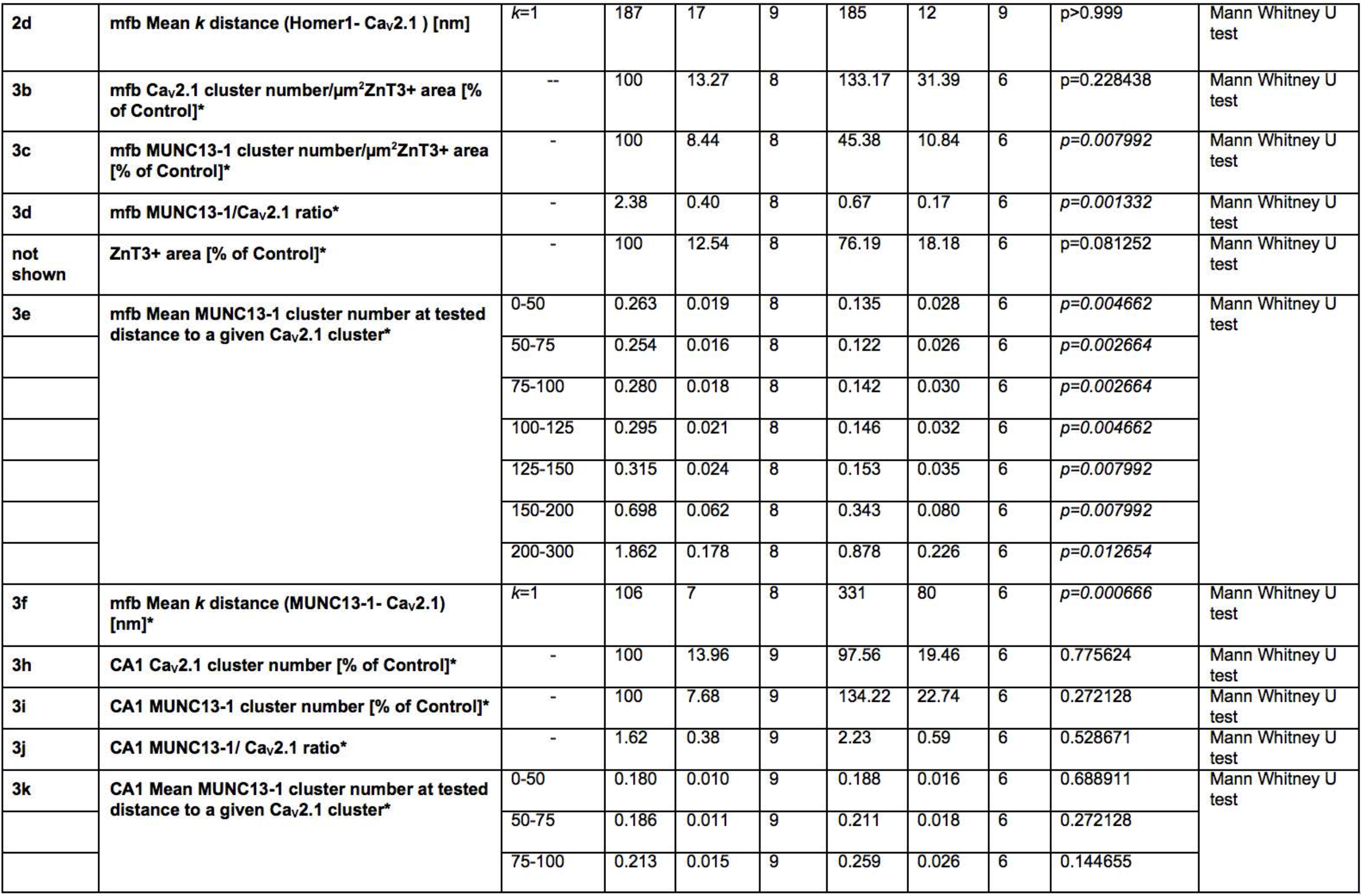
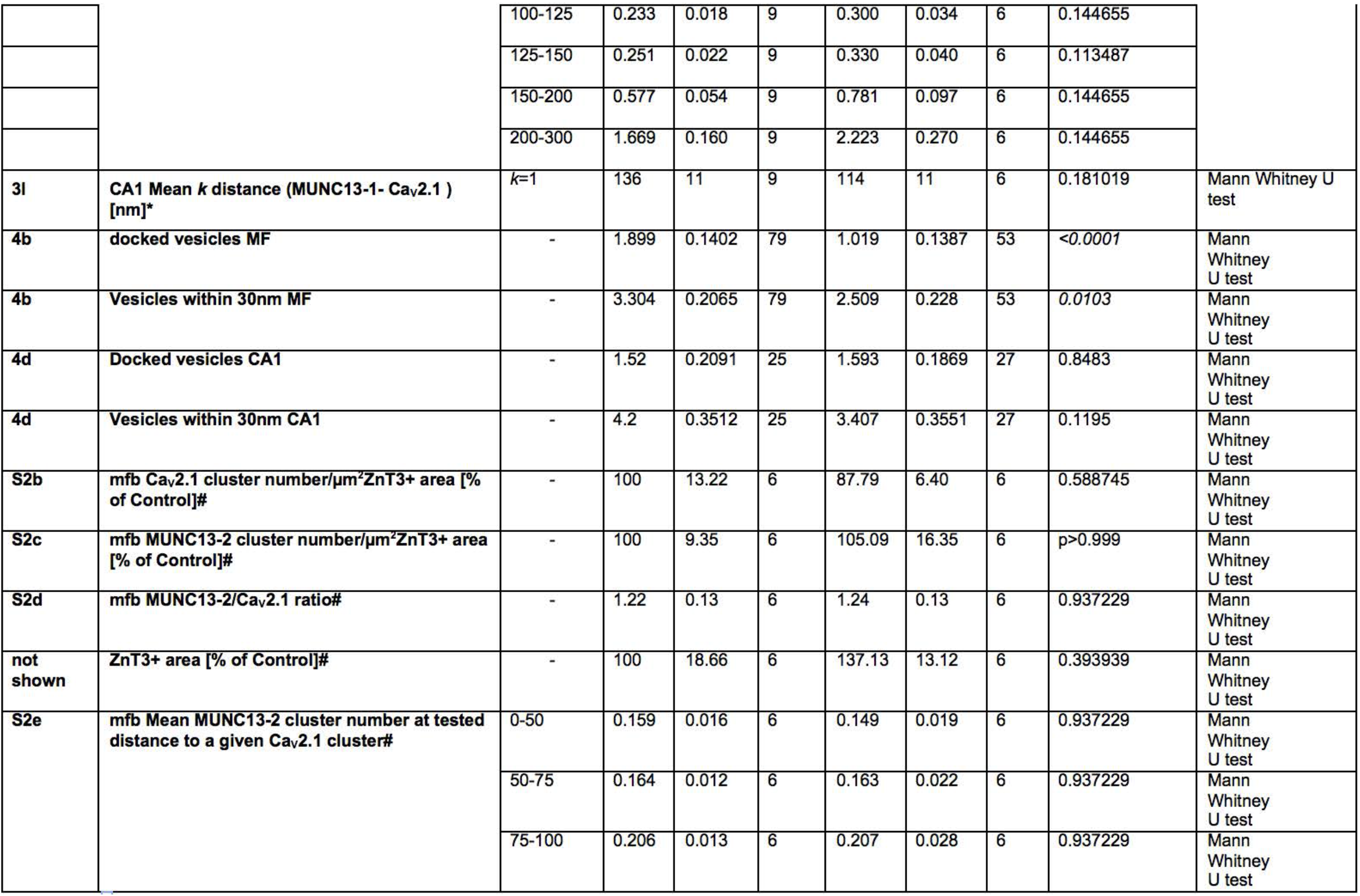
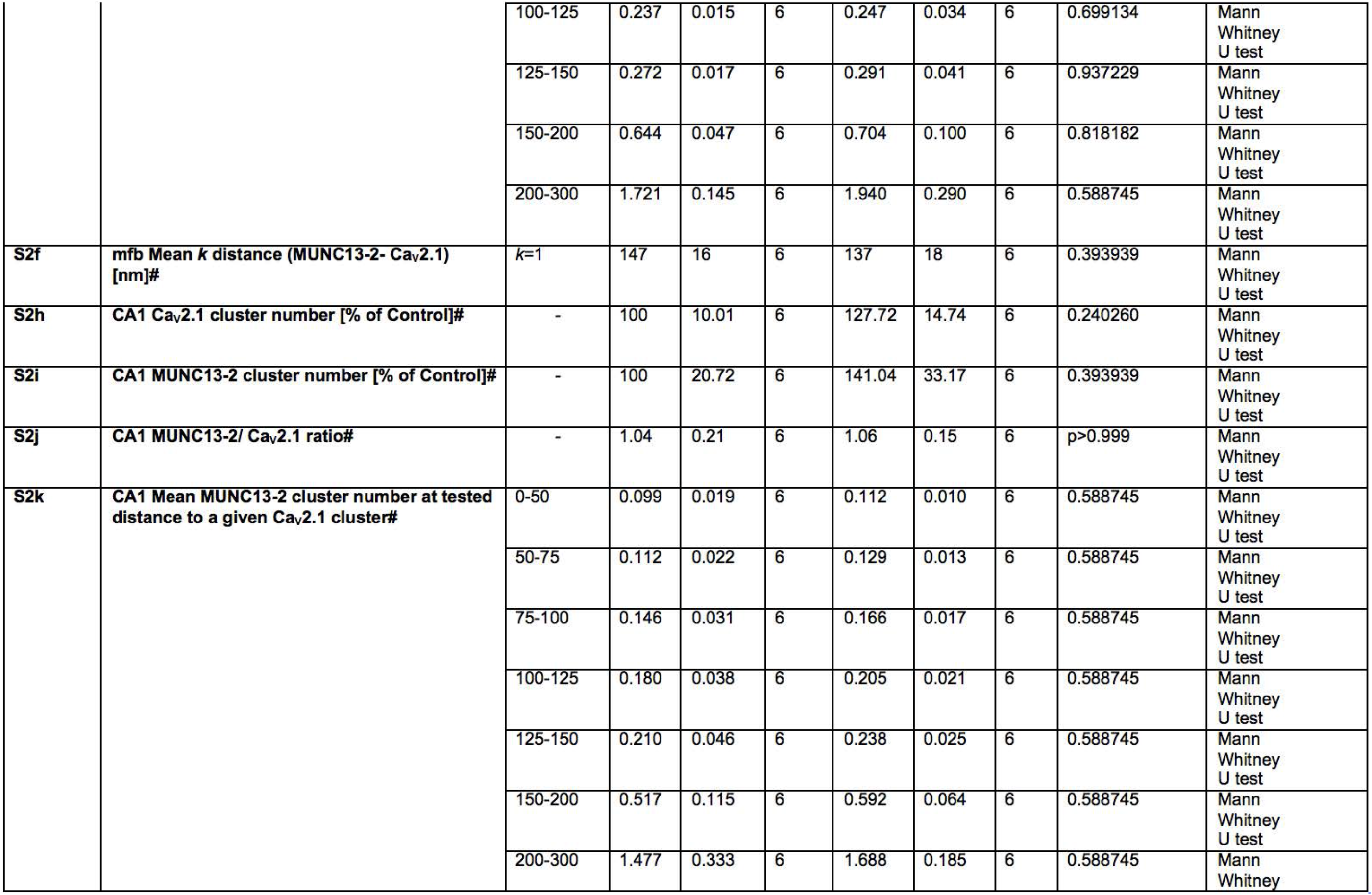
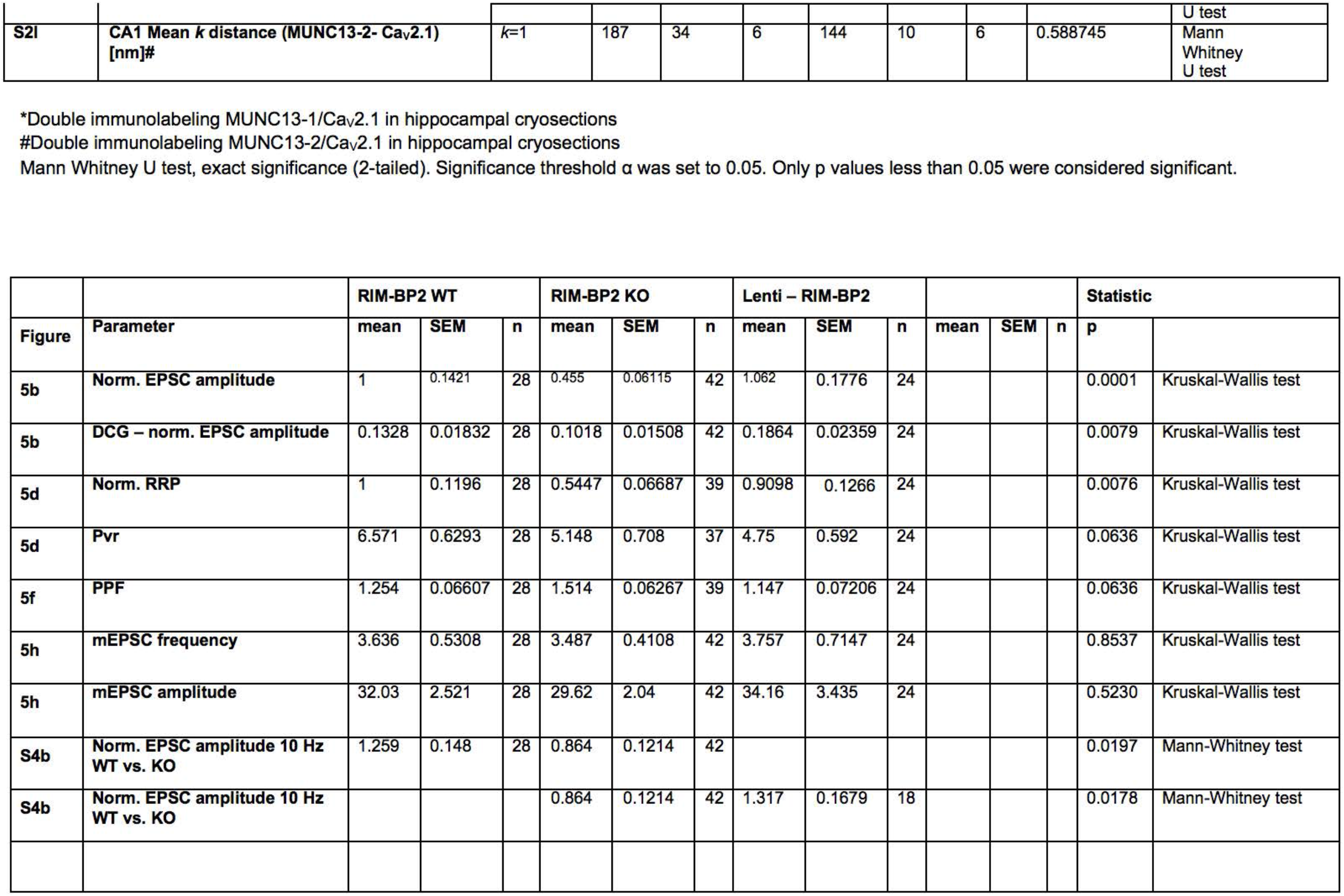
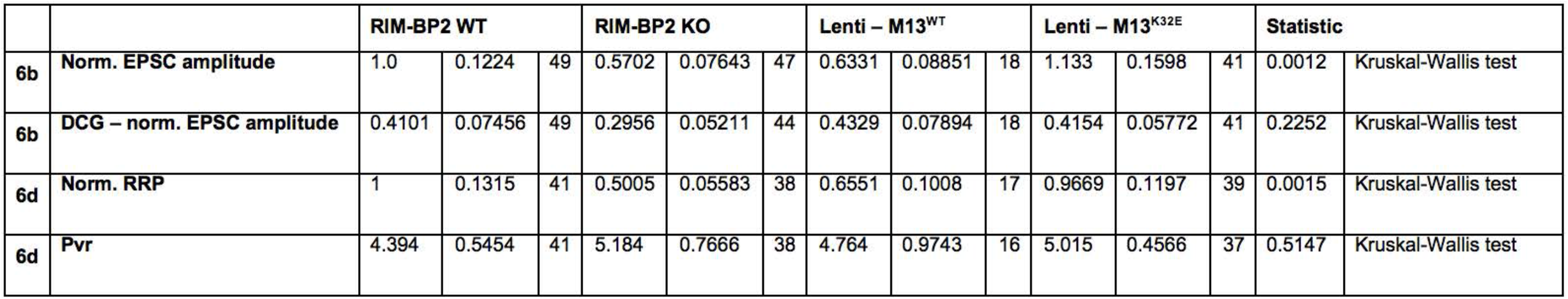
Summary of Statistical analysis.

